# Independent contributions of noradrenaline to behavioural flexibility and motivation

**DOI:** 10.1101/147512

**Authors:** Caroline Jahn, Sophie Gilardeau, Chiara Varazzani, Bastien Blain, Jerome Sallet, Mark Walton, Sebastien Bouret

## Abstract

Among neuromodulatory systems, the noradrenergic system remains one of the least understood. Several theories have pointed out its implication in behavioural flexibility and more recently in motivation, with a strong role in effort processing. Here, we designed a sequential cost/benefit decision task to test the causal role of noradrenaline in these two functions. We manipulated noradrenaline using clonidine, an alpha-2 noradrenergic receptor agonist, which reduces central noradrenaline levels. Clonidine had two distinct effects: it decreased choice volatility (without affecting the cost/benefit trade off) and reduced force production. Because the effects were independent, they cannot be accounted for by a non-specific effect on arousal. Altogether, these results support the global implication of noradrenaline in facing challenging situations in two complementary ways: by modulating behavioural volatility, which would facilitate adaptation depending on the lability of the environment, and by modulating the mobilization of resources to face immediate challenges.

## Introduction

Noradrenaline is among the most widespread neuromodulator in the brain and it has been involved in many cognitive processes, but its specific function remains unclear. The noradrenergic system was initially associated with vigilance and arousal (Kety, 1972; Harley, 1987; Aston-Jones et al., 1991; Berridge, 1991; Waterhouse et al., 1998; Berridge and Waterhouse, 2003), but several authors suggested that this role could extend to cognitive functions such as attention or learning and memory (Usher et al., 1999; Arnsten, 2000; Harley, 2007; Arnsten et al., 2012; Sara and Bouret, 2012; Sara, 2015). In particular, noradrenaline has been associated with various forms of cognitive flexibility (Deveauges and Sara, 1990; Bouret and Sara, 2004; Chamberlain et al., 2006; Tait et al., 2007; McGaugy et al., 2008; Guedj et al., 2016). Based on this work, several authors proposed original theories to capture the implication of the noradrenergic system in behavioural or cognitive flexibility (Yu and Dayan, 2005; Aston-Jones and Cohen, 2005; Bouret and Sara, 2005). More recently, we and others have also emphasized the potential role of noradrenaline in motivation, with a strong role in effort processing (Ventura et al., 2008; Bouret and Richmond, 2009; Zenon et al., 2014; Varazzani et al., 2015).

While aspects of these theories overlap, it has nonetheless been difficult to determine how to reconcile these different ideas as they have seldom been directly tested in the same experiment. Moreover, several recent studies have emphasized the strong relation between autonomic arousal and cognitive functions classically attributed to the noradrenergic system (Einhauser et al., 2008; Preuschoff et al., 2011; Nassar et al., 2012; Gee et al., 2017). This is based on the strong correlation between locus coeruleus (LC) firing and measures of autonomic arousal such as heart rate or pupil diameter, but this relation is far from being specific (Abercrombie & Jacobs, 1987; Joshi et al., 2016; Gee et al., 2017). Thus, one hypothesis would be that noradrenaline is simply associated with autonomic arousal and contributes to all processes linked with arousal in a highly non-specific fashion. In that frame, the various functions classically attributed to the LC (behavioural flexibility and motivation) could all be captured by a non-specific role in arousal and vigilance. Alternatively, noradrenaline would be involved in a set of specific and independent processes, which could not be captured by a generic function such as vigilance or arousal.

Therefore, the goal of these experiments is to test the causal role of noradrenaline in behavioural flexibility and motivation using a quantitative approach. To do this, we developed an original sequential decision making task, used computational modelling to identify precisely the cognitive processes of interest and then examined the consequences of manipulating central noradrenergic level using systemic injection of clonidine, an alpha-2 noradrenergic receptor agonist which decreases the firing of LC neurons and reduces central noradrenaline levels (Abercrombie and Jacobs 1987; Abercombie et al., 1988; Grant et al., 1988; Berridge and Abercrombie, 1999; Bouret and Richmond, 2009). To obtain a reward, a monkey should repeatedly squeeze a grip on a differing number of occasions to obtain rewards. The amount of force exerted on the grip was virtually not relevant for the task, since animals could complete a trial by exerting even a very small amount of force. The number of squeezes and the amount of reward that could be obtained on a given trial were instructed to the animal by the stimulus displayed on the screen. In most trials, after a number of responses on a given sequence, the animals were offered a choice to either complete the original sequence or change to execute a different one, with a different sequence length and reward size. This enabled us to look at the effect of clonidine on the cost/benefit trade-off during the choice about whether to persist or switch. In this setting, behavioural flexibility was assessed by measuring the variability in the choices. Motivation was assessed by measuring the willingness to work and the amount of force exerted on the grip, which was not a requirement for this task.

Clonidine had two distinct effects. First, in line with the role of noradrenaline in behavioural flexibility, clonidine dose-dependently decreased choice volatility: monkeys’ choices were more consistent under clonidine, but there was no effect on the cost-benefit trade-off. Second, in line with the role of noradrenaline in motivation and effort, clonidine dose-dependently reduced force production during the task. Because the effects on behavioural flexibility and motivation were statistically independent, they cannot be accounted for by a common confounding factor or a non-specific effect on arousal and vigilance. Altogether, these results support the global implication of noradrenaline in facing challenging situations in two complementary ways: i) by enhancing the mobilization of resources to face immediate challenges and ii) by increasing behavioural volatility, which would facilitate adaptation in a labile environment.

## Results

### Overview of the task

We trained three monkeys to perform the task depicted in figure 1. In each trial, monkeys performed series of actions (squeezing a grip 6, 8 or 10 times) to get a small, medium or large fluid reward (fig 1A). Note that the amount of force required to complete the trial was minimal and monkeys always succeeded to complete a squeeze when they tried. In 70% of trials, we introduced a choice by presenting an alternative option before monkeys could complete the trial. This alternative option, presented on the opposite side of the monitor compared to the current option, was also characterized by a given number of squeezes and a given reward size. Thus, monkeys could either choose to continue with the current option by squeezing the same grip as before, or switch to the other grip to start completing the alternative option and obtain the corresponding reward. As expected, the monkeys behavioural responses were affected both by the expected costs (number of remaining squeezes) and benefits (reward size), with no overall choice bias toward a specific side (left vs. right) or a specific option (current vs. alternative) (T-tests, p>0.05, see below). To capture the role of noradrenaline in behavioural flexibility and motivation, we then examined the influence of clonidine on corresponding behavioural measures in this task.

**Figure 1:**
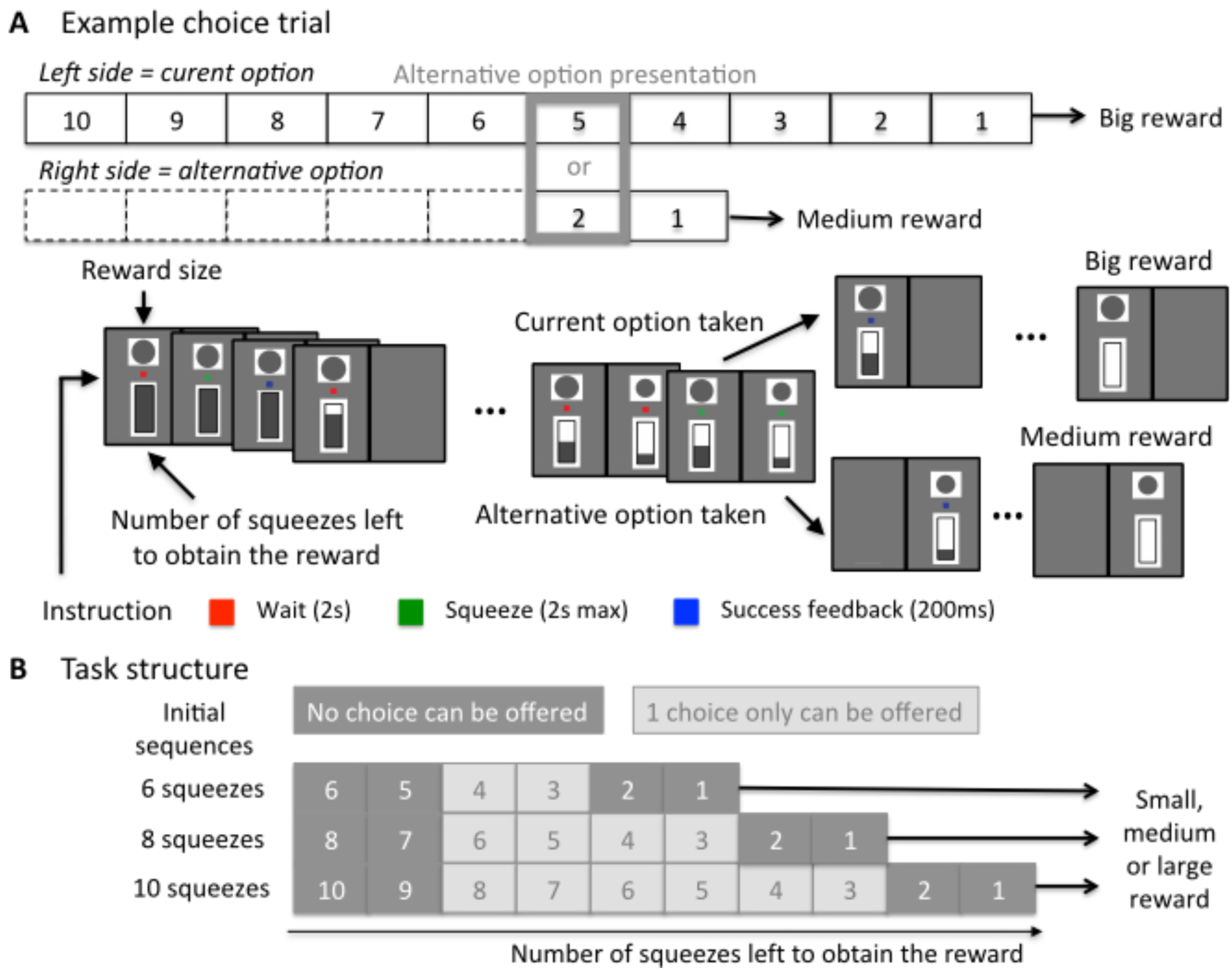
task. Monkeys performed an operant task where they have to exert a certain number (sequence) of squeezes on a grip to obtain fluid reward. They were sitting in a chair with two grips (right and left) facing a screen. The principle of the task is use the grip corresponding to the side where stimuli are displayed, wait when the dot is red, squeeze when it is green and a blue dot indicates a correct squeeze. All squeezes of a sequence must be performed correctly to obtain the reward. A squeeze is incorrect if monkeys do not squeeze above the minimum force threshold when the green dot is displayed, squeeze when the red dot is displayed, or use the wrong grip. After an incorrect squeeze, the same trial restarts. After all squeezes of a sequence are performed correctly, monkeys receive the fluid reward and a new trial starts. In 70% of trials, monkeys have the choice to continue with their current sequence by using the same grip or changing grip and perform an alternative sequence for an alternative reward size. *A) Example of a choice trial (70% of trials)*. The trial starts with the presentation of the option, which is defined by a side (here left), an initial sequence length (10 squeezes here, bottom cue) and a reward size (medium here, top cue). At each squeeze, the bottom cue indicates the remaining number of squeezes to perform (bottom cue). 5 squeezes before the end of the current, an alternative option is offered. Here, by squeezing the left grip, monkeys choose the current option and must perform the 5 remaining squeeze to obtain the megrim reward (top). By squeezing the right grip, monkeys choose the alternative option and must perform 2 squeezes to obtain the small reward (bottom). After all squeezes of a sequence are performed correctly, the gauge indicating the remaining number of squeezes in the sequence appears as empty and monkeys receive the fluid reward. After an inter-trial interval of 1 to 1.5s, another trial starts. B*) Task structure.* Initial sequences start with 6, 8 to 10 squeezes and lead to 3 sizes of reward (small, medium and big). In 30% of trials, no choice is offered and monkeys must perform the initial sequence to be rewarded. In 70% of trials one choice is offered during one of the squeeze in light grey (at least 3 trials after the beginning of a sequence and 3 trials before the end) on the figure.

### Effects of clonidine on behavioural flexibility: choices

We first measured the influence of clonidine on the average chosen number of squeezes and chosen reward size (fig 2A), which provides a global estimate of the animals’ relative sensitivity to costs and benefits. This measure was also not reliably affected by the treatment (linear regression taking into account subject variability, t(10)=1.85, p=0.09 and t(10)=0.65, p=0.53, respectively).

**Figure 2:**
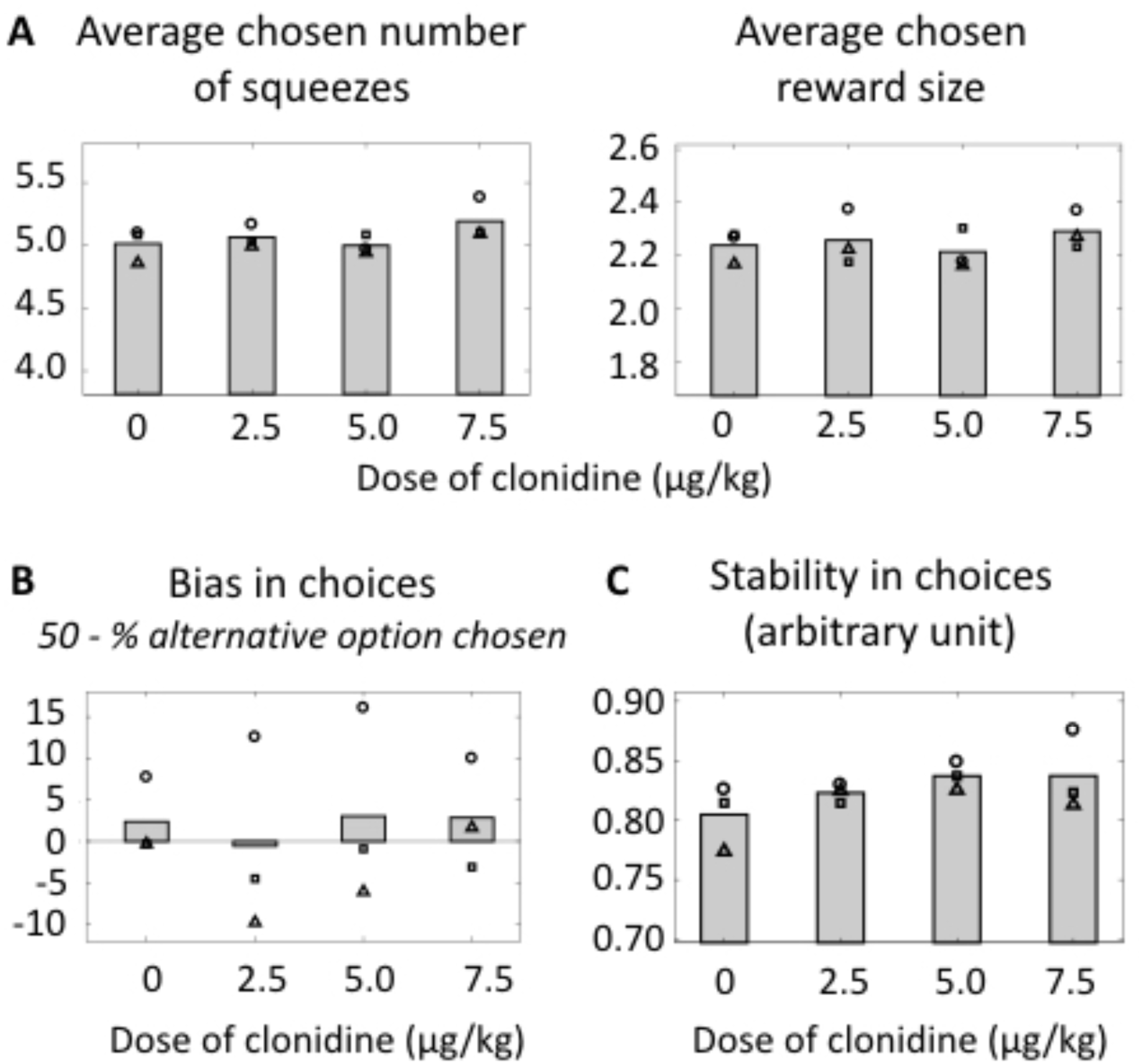
Effect of clonidine on decision-making. *A) Average chosen number of squeezes and reward size.* Average number of squeezes chosen to be performed at the choice point in each treatment condition. There was no effect of treatment condition. Size of average reward chosen (1 for small, 2 for medium and 3 for big) in each treatment condition. Same as A. There was no effect of treatment condition. Symbols correspond to each subject (circle: monkey A, square: monkey D, triangle: monkey E). There was no significant bias toward staying or switching across all treatment conditions.. *B) Overall bias for alternative or current option.* 50 - Mean across monkeys of the percentage of alternative option chosen for each treatment condition. Same as A. *C) Choices stability. Arbitrary unit.* Mean across monkeys and treatment condition. Same as A. Linear regression taking into account the variability across individuals revealed a significant positive linear effect of treatment condition (p<0.05).

We next examined the influence of clonidine on the choice bias towards the current versus alternative option, over and above the influence of expected costs (number of squeezes) and benefits (reward sizes). Because all options were offered in equal proportions as current and alternative options, any departure of the probability to take the alternative option from 50% would represent a bias toward staying or switching. At the group level, there was no significant bias toward staying or switching across all treatment conditions (linear regression taking into account subject variability, t(10)=0.52, p=0.52), and the bias was not different from zero in any condition (T-test, p>0.47, for all doses) (fig 2B).

We then looked the stability in choices across doses. As shown on figure 2C, there was a clear linear increase in stability across doses of clonidine, which means that with increasing doses of clonidine, the monkeys became increasingly likely to make the *same* decisions when faced with the *same* type of choice (linear regression taking into account variability across subjects, β=0.559±0.183, t(10)=3.06, p=0.01).

To capture the specific influence of clonidine on distinct components of decision-making, we built a simple choice model depicted in equations 1 and 2 in *Material and Methods*. In this model the value of each option corresponds to a trade-off between reward at stake and sequence length, controlled by a parameter k(sequence length). The probability to select a given option depends on (i) the value difference with the alternative and (ii) a fixed bias, e.g. a preference for either staying with the current option or taking the alternative, as well as (iii) the choice consistency, which determines the degree to which choices are consistent with the evaluation.

As shown on figure 3, we looked at the effect of the treatment on the three parameters of the choice model (k(sequence length), bias and consistency). The parameter k(sequence length) describing the relative sensitivity to reward and sequence length was significantly different from zero, indicating that monkeys readily integrated these two factors to guide their behaviour (all p<0.01). Had either the sensitivity to sequence length or reward size changed following administration of clonidine, this parameter would have varied For example an increase in effort sensitivity would have be translated in an increase in k(sequence length). But as shown on figure 3A, this parameter estimate was again not affected by the treatment (linear regression taking into account variability across subjects, t(10)=-0.10, p=0.92), indicating a lack of effect of clonidine on the cost-benefit analysis. In line with the previously described model-free analysis (fig 2B), there was no systematic bias to stay with the current option or switch to the alternative at the group level (bias parameter not significantly different from zero, all p>0.55) and no effect of treatment on this bias parameter (linear regression taking into account variability across subjects, t(10)=-0.10, p=0.92) (fig 3B). By contrast, clonidine induced a dose dependent increase in choice consistency (fig 3C). To analyse this formally, we ran a linear regression taking into account the variability across subjects. This revealed a significant linear effect of dose on the consistency parameter’s estimates (0=0.248±0.070, (10)=3.54, p<0.01).

**Figure 3:**
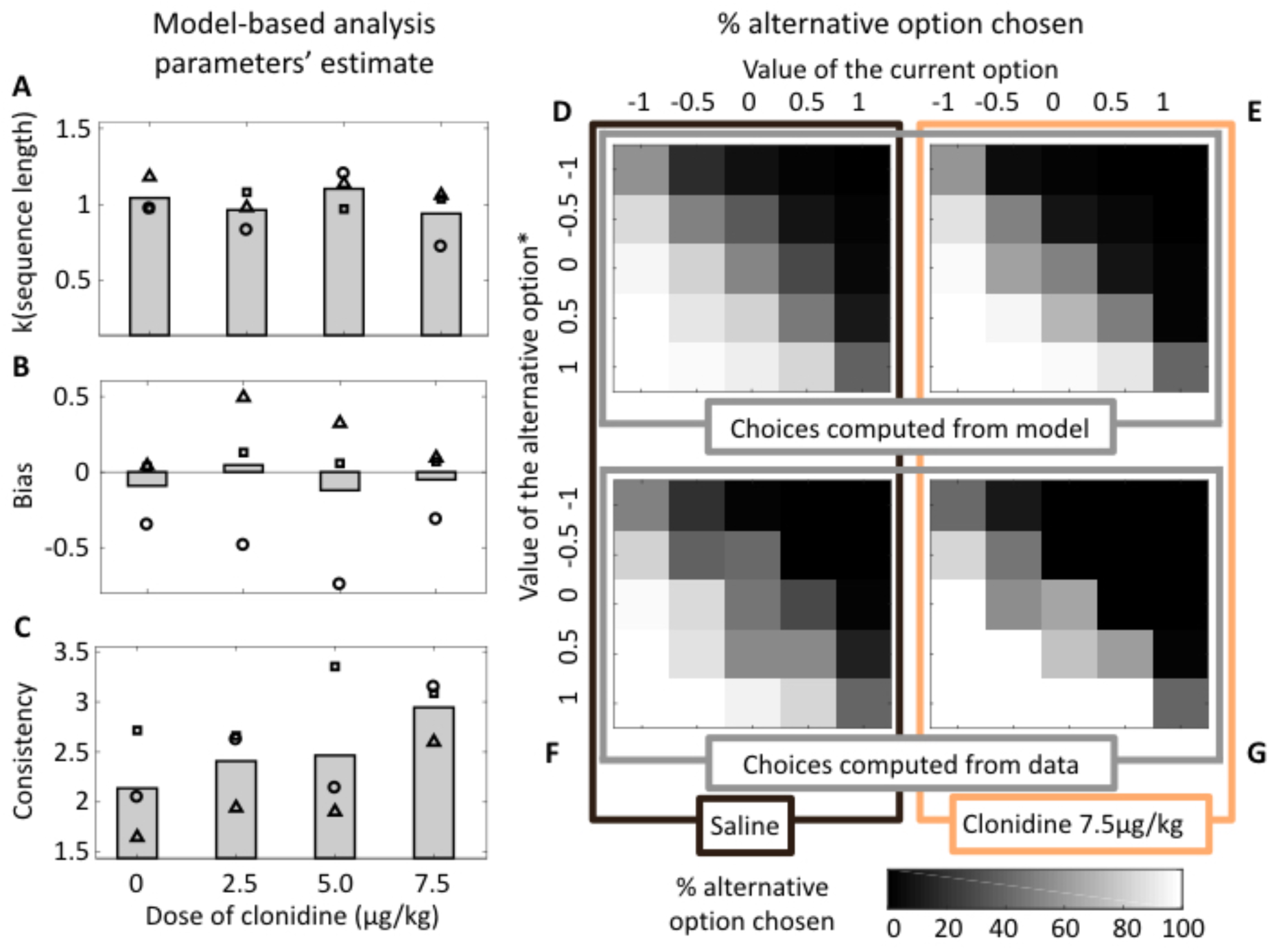
Clonidine specifically affects consistency in choice. *Mode-based analysis: parameters’ estimate. A) k(sequence length) parameter estimates.* Mean across monkeys of the k(sequence length) parameter estimates for each treatment condition in the choice model: Probability to take the alternative option = 1 /(1+exp(-(Value(alternative option)–Value(current option)+bias).consistency)) with Value(option) = Reward size – k(sequence length).Sequence Length, where sequence length is the remaining number of squeezes to perform to obtain the reward. Symbols correspond to each subject (circle: monkey A, square: monkey D, triangle: monkey E). There was no effect of treatment condition at the group level. *B) Bias parameter estimates*. Same as A. There was no effect of treatment condition at the group level. *C)* Same as A. Multi-level regression on estimated beta parameters taking into account the variability across subjects revealed a significant effect of treatment condition (p<0.01). *% alternative option chosen.* Mean across monkeys of the percentage of alternative option chosen depending on the value of the current option and the corrected value of the alternative option (V(alternative option)* = V(alternative option) + bias). D*) Model under saline.* Plot of the mean percentage of alternative option chosen computed with the choice model (estimates of the beta parameter). E*) Model under the higher dose of clonidine.* Same as D. F*) Data under saline.* Plot of the actual choices for the estimated values. G) *Data under the higher dose of clonidine.* Same as F.

Figure 3D-G illustrates the influence of the highest dose of clonidine on choices. The colour maps represent the proportion of alternative option chosen for different values of the current and the alternative options. Most stable choices are represented in white (subjects always choose the alternative option) and black (they always choose the current option). Grey colours represent less stable choices, with the maximum of randomness in along the diagonal, where the values of the two options are close. The more consistent the choices are, the smaller the area of this grey colours zone is. Figures 3C and D were generated using the estimated consistency parameter of the choice model under saline (fig 3C) and clonidine (fig 3E). Figures E and F with subjects’ actual choices. For both, under clonidine, the area occupied by stable choices expend at the expense of less stable choices, reflecting the increase in choice consistency following the pharmacological manipulation of noradrenaline.

### Effect of clonidine on reaction times

We next evaluated the effects of clonidine on reaction time across task conditions. We separated conditions where monkeys had to make a choice between two options from conditions where they only squeezed the grip to progress through the trial. As frequently observed, monkeys were slower to respond in choice than no-choice trials (fig 4A). We examined the influence of clonidine on reaction times in these two types of trials and a multi-level linear regression taking into account variability across subjects revealed a significant linear effect of choice (β=0.641±0.022, t(17)=6.30, p<0.001) and dose (β=0.070±0.007, t(17)=2.90, p<0.01), but no significant interaction (t(16)=0.13, p=0.89). Hence, clonidine significantly slowed down reaction times, but its effects were undistinguishable between choice and non-choice conditions. We also separated choice reaction times according to two levels of choice difficulty (depending on whether the dimensions to integrate to make the choice were congruent or not: *hard* and *easy choices*, respectively) and found a significant linear effect of choice difficulty (β=0.258±0.013, t(33)=7.52, p<0.001) and dose (β=0.083±0.008, t(33)=2.75, p<0.001), but once again no interaction (t(32)=-0.44, p=0.66). Both clonidine and choice difficulty increase reaction time but their effects are additive, indicating that clonidine does not interfere with the influence of difficulty on reactions times. Overall, monkeys’ reaction times were clearly modulated across conditions: animals slowed down when they had to make a choice, especially if it was difficult. High doses of clonidine also slowed down the monkeys but because its effects were equivalent across conditions (no interaction), it did not affect the behavioural effect of difficulty.

**Figure 4:**
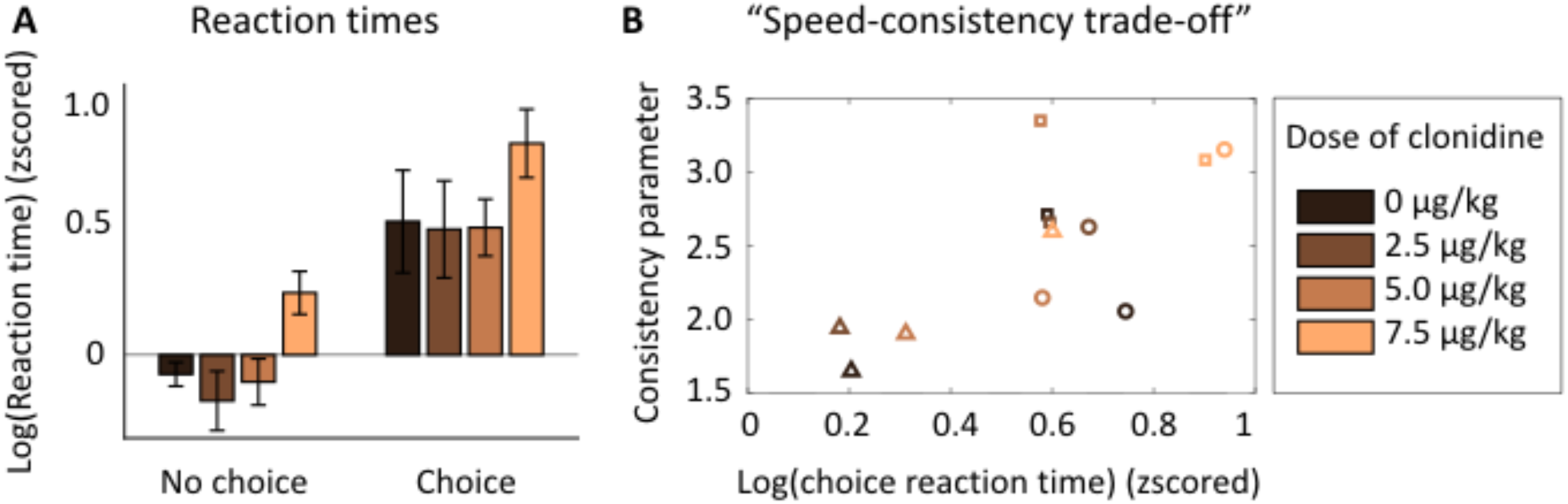
Effects of clonidine on reaction times. *A) Reaction times for no choice and choice.* Reaction times in 2 squeeze types (no choice and choice) for each treatment condition. No choice squeezes are matched to choice squeeze for position in the sequence and compared to choices in which subject did not change grip. Reaction time distributions for each grip and each monkey are logged z-scored (mean sets to zero and variance to one). Mean across monkeys, error bars represent standard errors to the mean. Colour code corresponds to treatment condition. Linear regression on log-transformed reaction times revealed a significant positive linear effect of choice (p<0.001) and treatment condition (p<0.01). *B) Correlation between choice reaction times and consistency parameter estimates.* Reaction times corresponds to the time between the display of the green dot and the crossing of the minimum force threshold for correct squeezes where monkeys had to make a choice and did not change grip. The consistency parameter was computed by fitting the choice model. They were computed for each subject and treatment condition. Same as A. The correlation between these two parameters was significant (p<0.01).

Together, our analyses therefore revealed two effects of clonidine on behaviour: it dose-dependently increased choice consistency and decreased choice reaction times. We examined the relation between these two effects across treatments and animals. We found a positive correlation (linear regression taking into account variability across subjects, β=0.321±0.095, t(10)=3.47, p<0.01) between the estimated consistency parameter and the choice reaction time (fig 4B). This correlation between the effect of treatments on reaction time and choice consistency suggests that clonidine affects a single functional entity, which we could refer to as “speed-consistency trade-off”. Altogether, this analysis provides a clear computational characterization of the contribution of the noradrenergic to behavioural flexibility in this task.

### Effect of clonidine on motivation: willingness to work

After assessing the implication of noradrenaline in behavioural flexibility, we examined the causal role of noradrenaline in motivation in this task. For that, we examined the influence of clonidine on two behavioural measures that are classically used to assess motivation, willingness to work and physical force production (fig 5). We measured monkeys’ willingness to work by counting the proportion of accepted squeezes. Since the action is very easy, monkeys never failed to complete a squeeze if they tried to, the number of squeezes that they accept to perform directly reflects their motivation to complete the trial. First, the willingness to work during 1- hour-long sessions was not significantly affected by dose (linear regression taking into account the variability across subjects, t(10)=-0.36, p=0.73) (fig 5A). Thus, clonidine did not have a global effect on the monkeys’ engagement in the task.

**Figure 5:**
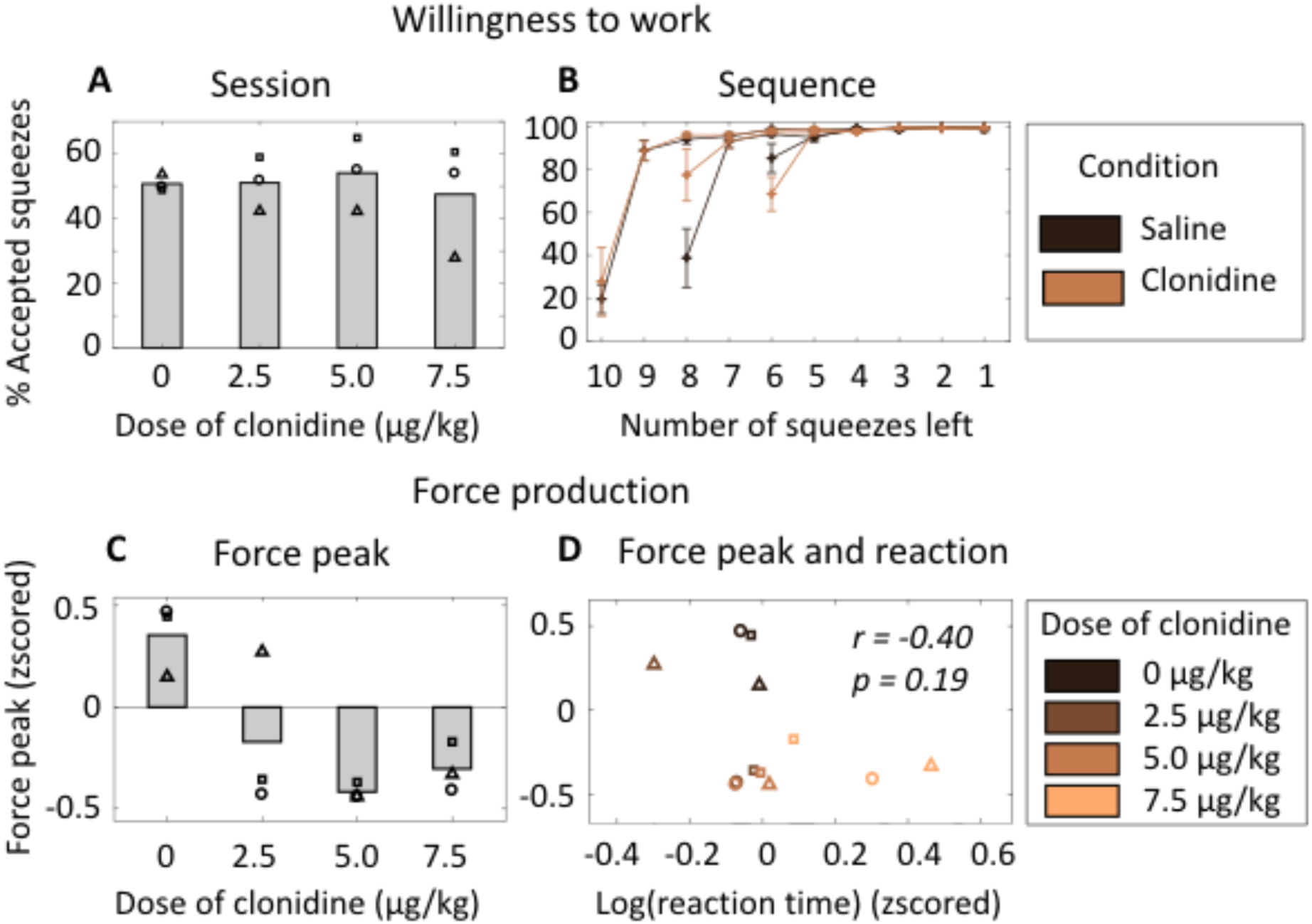
Effect of clonidine on motivation. *Willingness to work. A) Willingness to work during the session.* Number of accepted squeezes / total number of squeeze during the session (1 hour) for each treatment condition (in %). Mean across monkeys. Symbols correspond to each subject (circle: monkey A, square: monkey D, triangle: monkey E). There was no significant effect of treatment condition. *B) Willingness to work across the sequence.* Number of correct squeezes / total number of squeezes depending on the number of remaining squeezes to complete the sequence in trials where no choice was offered (in %). For simplicity, only the effect of treatment (colour code: saline vs. clonidine, all doses pooled) and sequence length are shown here. Mean across monkeys, error bars represent standard errors to the mean. *Force production. C) Force peak.* Maximal value of the force signal between two crossings of the minimal force threshold for correct squeezes. Force peak distributions for each grip and each monkey are z-scored (mean sets to zero and variance to one). Mean across monkeys. Same as A. Linear regression showed a negative effect of doses on peak force (p<0.01). *B) Peak force and reaction time*. Peak force is the same as in C. Reaction time corresponds to the time between the display of the green dot and the crossing of the minimum force threshold for correct squeezes for each treatment condition. Reaction time distributions for each grip and each monkey are logged and z-scored (mean sets to zero and variance to one). Symbols correspond to each subject (same as A). Colour code corresponds to treatment condition.

We also examined the effect of clonidine on the animals’ ability to adjust their behaviour across levels of progression through the sequences. We first examined willingness to work for the first squeeze of all sequences. We included every trial, since there was no way for monkeys to predict at the start of a sequence if a choice was going to be offered later in that sequence. We again examined the influence of reward size and sequence length on the willingness to perform the first squeeze using a linear regression taking into account the variability across subjects. This analysis revealed a significant negative effect of sequence length (i.e., the animals were less willing to engage on long sequences: β=-18.81±2.44, t(104)=-7.71, p<0.001) and a marginally significant positive effect of reward size (animals were more willing to engage for greater reward: 0=-4.79±2.44, t(104)=1.96, p=0.052), but no effect of dose of clonidine (t(104)=1.43, p = 0.16) and no interaction with sequence length (t(102)=0.20, p=0.84) or reward (t(102)=-0.65, p=0.52).

In a subsequent analysis, we examined the influence of clonidine on the adjustments of willingness to work across the steps of a trial, as a function of the upcoming sequence length and reward size (fig 5B). Indeed, monkeys displayed a sharp increase in their willingness to work after the first squeeze, which can be related to the engagement in the sequence. We fitted the curves depicted in figure 5B with equation 3 for all trials during which no choice was offered. In this model, k(intercept) controls the intercept (initial willingness to work) and k(slope) the slope of the rise of the willingness to work across the sequence. We then examined the influence of reward and sequence length on each of these parameter estimates using a multi-level linear regression taking into account the variability across monkeys. The parameter k(intercept) displayed a significant positive linear effect of sequence length (β=0.160±0.019, t(104)=8.09, p<0.001) and a negative effect of reward size (β=- 0.048±0.019, t(104)=-2.44p<0.05), but again there was no significant linear effect of dose (t(103)=-1.85, p=0.07), and no interaction with either the sequence length (t(101)=0.32, p=0.75) or the reward size(t(101)=-0.12, p=0.91). By contrast, neither the task factors (reward and sequence length) nor the dose of clonidine and its interaction with the tasks factors affected the parameter k(slope), which captured the slope of the change in willingness to work across the sequence (all p > 0.30).

In short, monkeys displayed robust adjustments of their willingness to produce the action across conditions, defined by the distance to reward and the amount of expected reward at the end of the sequence, but this was unaffected by clonidine. This implies that clonidine did not affect the monkeys’ general willingness to work, ruling out a non-specific effect on arousal.

### Effect of clonidine on motivation: Force production

Lastly, we examined the effect of clonidine on another key component of motivation: force production. As shown on figure 4C, force peak was significantly decreased under clonidine treatment (linear regression, t(10)=-3.24, p<0.01). Moreover, as described in an earlier section, clonidine increased overall the reaction times (linear regression, t(10)=2.63, p=0.02). Hence we considered the possibility that clonidine had a global, non-specific effect on arousal or vigilance, which would be responsible for both longer reaction times and smaller force peaks. Such a scenario implies a strong relation between the effects of clonidine on force peak and reaction time. We first compared the effect on force peak and reaction time in a 2-way ANOVA with factors dose and measure (force peak vs. reaction time). We found a significant main effect of dose (F(3, 16)=5.25, p=0.01), measure (F(1,16)=4.72, p=0.04), but importantly also a significant interaction between the two (F(3, 16)=8.59, p=0.001), indicating that the effect of clonidine differs significantly between the two measures. We furthered this analysis using a linear regression (fig 4D): there was no reliable correlation between the two measures (r=-0.4, p=0.19). Moreover, we ran separate linear regressions without the highest dose, we found that the effect on force peak was still significant (t(7)=-3.93, p=0.005) whereas it was not the case for reaction times (t(7)=0.16, p=0.88), implying that the linear effect was mostly due to the last dose. We also found a greater linear effect of dose on (β=-0.222±0.068) than on reaction times (β=0.106±0.040). Overall, the analyses imply that clonidine had a greater impact of effort production than willingness to work and reaction times.

Finally we found no significant correlation between the consistency parameter and the force peak, (t(10)=-1.39, p=0.15). This shows that the effect of clonidine on force production and decision-making (“speed-consistency trade-off”) are independent and probably not due to a global effect on arousal/vigilance.

## Discussion

In the present work, we used a novel decision-making task, which required monkeys to make sequential actions for reward and, on a majority of trials, to choose whether to stick with the current sequence or to switch to an alternative based on the costs and benefits of the options. This task allowed us to independently evaluate two classes of functions thought to involve noradrenaline: behavioural flexibility and motivation. Using systemic injections of clonidine, at doses that specifically decrease the level of noradrenaline in the brain (Kawahara et al., 1999; Fernandez-Pastor et al., 2005), we showed that noradrenaline was causally involved both in specific aspects of behavioural flexibility (speed consistency trade-off) and also in motivation (willingness to work and force production). Importantly, the effects on force were specific to this parameter and did not relate to other measures of motivation such as response speed or willingness to work, negating any general changes in arousal or vigilance.

### Noradrenaline regulates behavioural flexibility: speed/consistency trade-off

The first major effect of clonidine treatment was on choice consistency, thus extending the causal role of noradrenaline to value-based decision-making. Model-free analysis showed that clonidine induced a decrease in choice variability. In other words, when given clonidine, the monkeys became more likely to repeat the same choice when presented with particular pairs of choice options. This was also backed up by our model-based analysis, where we separated the evaluation from the option selection components in a simple “value-first” model (Padoa-Schiopa, 2011). These two processes were controlled by 3 distinct parameters: *k(sequence length)* for the cost-benefit evaluation, a *bias* and *consistency* for the option selection. This model-based approach linked the behavioural change induced by clonidine specifically with an increase of the consistency parameter, independent of the valuation process and the bias parameter. Hence this effect was due to a decrease in choice variability rather than to a systematic biasing of the choices in any direction (such as by reward, cost, side or current vs. alternative option).

Variability in choices is assumed in many models of decision-making, but its functional role remains debated. It has been proposed that it arises from random noise driven by internal neural variability (Wang, 2002; Faisal et al., 2008; Drugowitsch et al., 2016) and in that case, noradrenaline would control the amount of noise that is allowed in the computation (Aston-Jones and Cohen, 2005). Along those lines, it has recently been proposed that noradrenaline controls the precision of sensory cortical representation (Warren et al., 2015).

Optimal economic theory stipulates that the behaviour is optimal when there is no noise, meaning that the choices follow exactly the values of the options. In that sense, clonidine would seem to make monkeys more optimal since it increases choice consistency. But this absence of noise is only optimal in a constant environment. In a more uncertain and dynamic environment, however, noise in choices is thought to facilitate adaptation (Aston-Jones and Cohen, 2005; Yu and Dayan 2005; Nassar et al., 2012; Wilson et al., 2014). Thus, an increase in choice consistency under clonidine might also be interpreted as a decrease in efficacy if placed in a labile and/or uncertain environment. Importantly, our experiment shows that an effect of noradrenaline manipulation on choice variability can be observed even without a change in reward rate. This implies that the action of noradrenaline on choice consistency is systematic and generic, rather than dependent upon the specific task contingency. As such, these results are consistent with two recent studies. The first showed that specifically enhancing LC inputs to rat anterior cingulate cortex triggers behavioural variation (Tervo et al., 2014). The second showed that systemic clonidine increases decisiveness in rats by reducing the deliberative search process and representation of the unchosen path in the hippocampus in a spatial decision-making task (Amemiya and Redish, 2016).

Our work also indicates that the effect of clonidine on choice consistency was associated with a slowing of choice reaction times, in line with the idea that noradrenaline affects an internal decision variable, a specific function that affects both consistency and speed. The effect on speed consistency trade-off resonates with the idea that efficient deliberation takes time, such that slow decisions are also more reliable because they are based on a more thorough evaluation of subjective costs and benefits (Jocham et al., 2014).

Thus, not only is our work further supporting the implication of the noradrenergic system in behavioural flexibility, but it provides a clear computational characterization of its role. Noradrenaline promotes behavioural volatility by favouring faster and less reliable choices, thereby inducing a more noisy behaviour, and without affecting the cost-benefit trade-off.

### Noradrenaline regulates motivation: force production

In line with the conclusions of our recent electrophysiological studies (Varazzani et al., 2015), these experiments using direct manipulation of the noradrenergic system demonstrated its causal implication in motivation. Clonidine dose-dependently and specifically reduced the amount of force produced, and this effect on force was independent of the effects on reaction time, ruling out a simple interpretation in terms global motor impairment. It is also unlikely to be caused by a global effect on arousal or vigilance, as clonidine had no impact on either the animals’ willingness to work or on their ability to switch from the current to the alternative option. The influence of clonidine on force production was independent of task conditions, including reward. Indeed, the amount of force produced was not contingent in this task and the effect of clonidine was equivalent across reward conditions. Hence clonidine did not affect incentive processes, as it is often the case with dopaminergic treatments (Denk et al., 2005; Lebouc et al., 2016; Yohn et al., 2016; Zenon et al., 2016).

Thus, the effect of clonidine causing reduced force production seems to be relatively specific to the action requirements, but did not affect the overall cost-benefit analysis. Indeed, neither the initial choice to engage with the sequence, nor the binary choice in the middle of the sequence to stick with the initial option or to switch to the novel alternative were affected by the treatment as shown in the model-based analysis. At first, this might appear surprising since squeezing the grip multiple times could be taken as an effort. But since the minimal force to validate a squeeze was very small and monkeys always succeeded to reach it if they initiated the action, physical effort is unlikely to be a major component of the cost in this task. Moreover, previous studies using a similar task suggest that some monkeys could treat sequences as delay, and neglect the motor cost relative to the temporal discounting effect of the sequence (Minamimoto et al., 2012). Given the small number of animals in monkey studies, this effect remains difficult to evaluate. Finally, this is in line with recent electrophysiological data in an effort-reward trade-off task (Varazzani et al., 2015) showing an activation of LC neurons correlated with the amount of force produced on a grip at the time of executed action, but not when evaluating this option. In other words, the LC neurons only encoded the effort component at the time of when monkeys needed to actually mobilize energy to produce the action and not when choosing whether to act in the first place. This reinforces the idea that noradrenaline plays a specific role in actually producing the effort – mobilizing energy to face a challenge – as we suggested earlier (Bouret and Richmond, 2015; Varrazani et al., 2015). It is intriguing that such a role is complementary yet distinct from the influence of the other major catecholamine, dopamine, which is known to be key for assessing the value of working through sequences of actions for reward but is perhaps not required to overcome force constraints (Ishiwari et al., 2004; Gan et al., 2010; Pasquereau and Turner, 2013; Varazzani et al., 2015; Salamone et al., 2016). A key question for future studies will be to directly contrast the precise roles these neurotransmitters play in effort-based decision-making.

Last, we found that the effects on force production were not correlated with choice consistency. Hence the two effects were independent, further ruling out an interpretation in terms of global, low level process such as arousal or vigilance. This is probably due to the action of clonidine on different networks. It has recently been shown different populations of LC neurons project to the prefrontal cortex and the motor cortex (Chandler et al., 2014). These two distinct networks could underlie the effects on choice consistency and force production respectively. But irrespectively of the underlying neurobiological mechanisms, this work demonstrates that these two facets of noradrenergic functions are relatively independent, and specific. In other words, the implication of noradrenaline in cognition and behavior cannot be reduced to arousal or vigilance, even if LC activity strongly correlates with autonomic arousal. One of the upcoming challenges will be to understand the neuronal mechanisms underlying these specific operations, and how they articulate with functions of other neuromodulatory systems such as dopamine and serotonine.

### Conclusion

To conclude, these results delineate the causal implication of noradrenaline in behavioural flexibility and motivation. As only particular behavioural functions were altered by clonidine, we can go beyond an interpretation based on a global arousing effect of noradrenaline. Instead, our results are compatible with the idea that noradrenaline is involved in facing challenges through two specific and complementary actions: i) an increase in behavioural volatility and ii) the mobilization of physical resources to face immediate challenges. This proposal relies on the assumption that these two processes are adaptive to solve most challenges. While the former would facilitate adaptation in an uncertain or changing environment, the later is clearly advantageous in an environment where you must compete for resources and achieve the goals that are set. Key challenges remain to understand how this system articulates with other neuromodulators in adaptive behaviours and what is its precise action on its target networks.

## Materials and Methods

### Monkeys

Two male rhesus monkeys (Monkey A, 15 kg, 5 years old; Monkey D, 15 kg, 6 years old) and one female (Monkey E, 4.5 kg, 3 years old) were used for the experiment. Their access to water was restricted and during testing days (Monday to Friday), they received water as reward. All experimental procedures were designed in association with the Institut du Cerveau et de la Moelle Epiniere (ICM) veterinarians, approved by the Regional Ethical Committee for Animal Experiment (CREEA IDF no. 3) and performed in compliance with the European Community Council Directives (86/609/EEC).

### Task

Each monkey sat in a primate chair positioned in front of a monitor on which visual stimuli were displayed. Two electronic grips (M2E Unimecanique, Paris, France) were mounted on the chair at the level of the monkey’s hands. Monkeys were not constrained to use one hand or the other to squeeze the grips. Each grip corresponded to one side of the screen. Water rewards were delivered from a tube positioned between the monkey’s lips. Behavioural paradigm was controlled using the REX system (NIH, MD, USA) and Presentation software (Neurobehavioral systems, Inc, CA, USA).

The task consisted of performing sequences of squeezes on a grip to obtain rewards. At the beginning of each trial, the length of the sequence (number of squeezes) and the size of the reward were indicated by two different cues that appeared simultaneously with a red dot on either the left or right side of the screen (counterbalanced across trials) (Fig 1). There were nine initial options defined by three initial sequence lengths (6, 8 and 10 squeezes) and three reward sizes (small, medium and big). After a fixed delay of 2s, the red dot turned green and to initiate a trial, monkeys had 2s to perform a squeeze above the minimum force threshold with the grip corresponding to the side of the screen where stimuli were displayed. The threshold was manually calibrated during the training phase, so that monkeys would always reach it if they squeezed the grip (bell-shaped force profile). After a correct squeeze, the dot turned blue for 200ms. Then, the dot turned red again and the cue corresponding the sequence length changed to show the number of remaining squeezes to complete the sequence. After an incorrect squeeze, the stimuli disappear and the same trial restarted from the beginning of the sequence after 1 - 1.5s of inter-trial interval delay. A squeeze was incorrect if monkeys squeezed the wrong grip, squeezed any grip when the dot was red or did not reach the minimum force threshold 2s after the dot turned green. After the last correct squeeze of the sequence, the dot disappeared, the cue indicating the number of remaining squeezes was at zero and monkeys received the size of the reward corresponding to the reward cue. At the end of the reward delivery, a new trial started after 1 - 1.5s of inter-trial interval delay.

In 30% of trials, monkeys had no option other than to complete the initial sequence. However, in 70% of trials, monkeys were given the choice during the sequence to take an alternative option (Fig 1A). The alternative option was presented on the opposite side of the screen and occurred at least three squeezes after the beginning the initial sequence and at most three squeezes before the end of it (fig 1B). To choose this option, monkeys had to switch to squeezing the corresponding grip when the dot turned green. Only one alternative option was offered per trial and it was presented only once during the sequence. In 10% of trials, the alternative option was the *same* as the current option (i.e., same reward size and same remaining sequence length). In 20% of trials, the two dimensions of the choice were *congruent*: the alternative option had either a longer / same sequence length and a smaller / same reward size or a shorter / same sequence length and a larger / same reward. In the remaining 40% of trials, the two dimensions of the choice were *incongruent*: the alternative option either had a longer sequence length but a bigger reward size or a shorter sequence length but a smaller reward. The sequence length and reward size of the alternative option were drawn so that: i) if option A was offered as a current option and B as alternative on one trial, it was equally probable that A would be offered as a alternative and B as a current option on another trial, ii) all sequence lengths (3 to 8 squeezes) were equally probable for the current and the alternative options, and iii) before the choice, the numbers of squeezes performed were counterbalanced across sequences. As a consequence, starting with a long sequence was more probable (11 out of 18 trials) than a medium (5 out of 11 trials) and a short (2 out of 11 trials) sequence. All reward sizes and sides were equally probable.

### Pharmacological procedure

We used three doses of clonidine - 2.5, 5.0 and 7.5μg/kg - which is a selective alpha-2 noradrenergic receptor agonist that suppresses LC firing and consequently noradrenaline release at the doses that we used (Kawahara et al., 1999; Fernandez-Pastor et al., 2005). The doses that we used were below the sedative effect threshold determined in rats (Sara et al., 1995; Lapiz and Morilak, 2006) and monkeys (Bouret and Richmond, 2009), but elicited a subjective feeling of sedation humans (Jäläkä et al., 1999). Clonidine hydrochloride (C7897, Sigma-Aldrich, St. Louis, MO, USA) solutions were prepared freshly each day by dissolution in 1mL saline for monkey A and D, and 0.5mL for monkey E. The same volume of saline solution was given in saline condition. Drug or saline solution was injected intramuscularly 20min before testing in monkeys’ home cages. Each dose or vehicle was given for five consecutive days (Monday to Friday). Order of drug and saline weeks (one drug week per dose and 2 saline weeks) was randomly assigned for each animal.

### Data analysis

Data were analysed with Matlab software (MathWorks). To assess changes in the decision process, we looked at three variables: (i) the proportion of alternative options chosen per session, (ii) average chosen number of squeezes, (iii) the average chosen reward size and (iv) the stability in choices. The stability was computed by assessing for each possible combination of differences in reward size (5 possibilities: -2, -1, 0, +1, +2) and sequence length (5 possibilities: -4, -2, 0, +2, +4) between the current and the alternative option if the same choice (taking the alternative or the current option) was made. We then took the average for these 25 possibilities. The stability is therefore maximal if monkeys are perfectly stable in there choices and minimal if there are completely random.

We also built a simple decision model where the value of each option (the current and the alternative) is computed as:

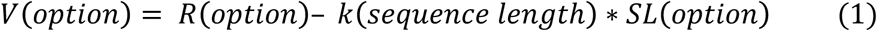

where V(option) is the value, R(option) the reward size and SL(option) the sequence length of the considered option. Sequence Length corresponds to the remaining number of squeezes to perform to obtain the reward. The values of the two options are compared to determine the probability to take the alternative option as follows:

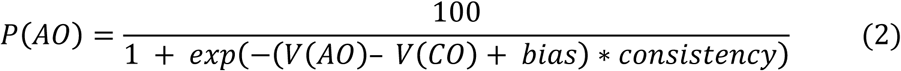

where P(AO) is the probability to take the alternative option, V(AO) and V(CO) the values of the alternative and current options respectively computed with equation 1. The three parameters k(sequence length), bias and consistency in equations 1 and 2 were estimated by inverting the model so as to minimize the free energy, using a variational Bayes approach under Laplace approximation (Friston et al., 2007; Daunizeau et al., 2009), implemented in a Matlab toolbox (available at http://mbb-team.github.io/VBA-toolbox/; Daunizeau et al., 2014).

Reaction time corresponds to the time between the display of the green dot and the crossing of the minimum force threshold for correct squeezes. Reaction time distributions for each grip and each monkey log-transformed and z-scored (mean set to zero and variance to one). We compared reaction times across different squeeze types. When comparing *Choice* and *No Choice* reaction times, No Choice reaction time corresponds to trials where no choice was offered but in principle could have been (fig 1B) and Choice reaction time corresponds to points in the sequence where monkeys were presented with an alternative option but stayed with the original option. When looking at the effect of difficulty on choice reaction times, *Easy* choices correspond to ones where the two dimensions of the choice are strictly congruent and monkeys do not change grip. *Hard* choice squeezes correspond to squeezes where the two dimensions of the choice are strictly incongruent and monkeys do not change grip.

Motivational changes with treatment were assessed by two variables: force peak and willingness to work.

To calculate force peak, force time series for both grips were low-pass filtered at 15 Hz (zero-phase second-order Butterworth filter) and we took the maximal value of the force signal between two crossings of the minimal force threshold.

Willingness to work corresponds to the proportion of accepted squeezes per session. Willingness to work at the beginning of each sequence was computed by taking the first squeeze of all trials. Willingness to work across sequences of given length and reward size were estimated by taking all trials where no choice was offered in a given drug condition for each monkey and fitted using the following model:

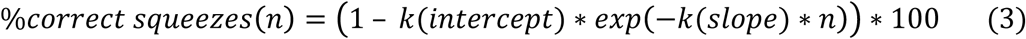

Where n is the number of squeezes done in the sequence. Parameters k(intercept) and k(slope) were estimated for each dose and each monkey using the same procedure as for the parameters of equations 1 and 2.

### Statistical Analysis

Data are plotted as mean +/− standard error to the mean. Statistics used are indicated in the *Results* sections. Comparisons between means were performed using parametric tests (ANOVA and T-test). We performed linear regression on z-scored distributions (reaction times and force peaks) using the function glmfit in Matlab. In cases when distributions were not z-scored, we fitted an intercept for each subject hence taking into account the variability in mean across subjects using the function fitlme in Matlab. The general equation was:

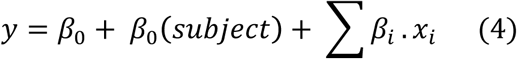

where y is the data, *β*_0_ a constant, *β*_0_ subject a constant fitted for each subject, *x_i_* the experimental factors and *β_i_* their weights in the linear regression. T-tests were performed on weights distributions. In all cases t-values and degrees of freedom are given according to statistical analysis reports of glmfit and fitlme. All statistical tests were two-sided. *P* > 0.05 was considered to be not statistically significant. We evaluated the quality of our models’ fit using balanced accuracy (between 0 and 1) computed as:

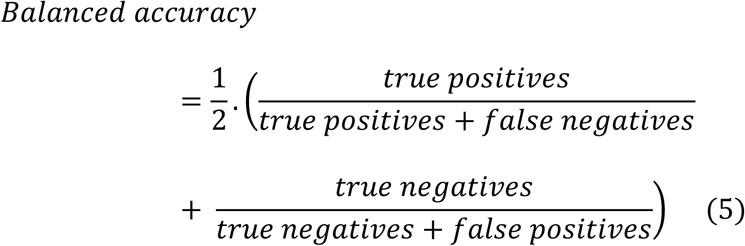

which was between 0.80 and 0.86 for all fits (Brodersen et al., 2010).

## Acknowledgments

We thank Morgan Weissenburger and the animal housing facility staff for their assistance and care of the animals. We also thank Nicolas Borderies for discussion. This research was funded by the ERC BIOMOTIV, the Paris Descartes University doctoral and mobility grants and the Wellcome Trust fellowships (MEW: 090051 and 202831 /Z/16/Z, JS: WT1005651 MA).

